# Redox regulation of yeast Hsp70 modulates protein quality control while directly triggering an Hsf1-dependent cytoprotective response

**DOI:** 10.1101/2022.04.08.487668

**Authors:** Alec Santiago, Kevin A. Morano

**Affiliations:** Department of Microbiology and Molecular Genetics, McGovern Medical School at UTHealth Houston, Houston, TX USA; MD Anderson UTHealth Graduate School at UTHealth Houston, Houston, TX USA

**Keywords:** Hsp70, chaperone, redox, cysteine, thiol modification, reactive oxygen species, proteostasis

## Abstract

Neurodegenerative disease affects millions of Americans every year, through diagnoses such as Alzheimer’s, Parkinson’s, and Huntington’s diseases. One factor linked to formation of these aggregates is damage sustained to proteins by oxidative stress. Cellular protein homeostasis (proteostasis) relies on the ubiquitous Hsp70 chaperone family. Hsp70 activity has been previously shown to be modulated by modification of two key cysteines in the ATPase domain by oxidizing or thiol-modifying compounds. To investigate the biological consequences of cysteine modification on the Hsp70 Ssa1 in budding yeast, we generated cysteine null (cysteine to serine) and oxidomimetic (cysteine to aspartic acid) mutant variants of both C264 and C303 and demonstrate reduced ATP binding, hydrolysis and protein folding properties in both the oxidomimetic as well as hydrogen peroxide-treated Ssa1. In contrast, cysteine nullification rendered Ssa1 insensitive to oxidative inhibition. The oxidomimetic *ssa1-2CD* (C264D, C303D) allele was unable to function as the sole Ssa1 isoform in yeast cells and also exhibited dominant negative effects on cell growth and viability. Ssa1 binds to and represses Hsf1, the major transcription factor controlling the heat shock response, and the oxidomimetic Ssa1 failed to stably interact with Hsf1, resulting in constitutive activation of the heat shock response. Consistent with the in vitro findings, *ssa1-2CD* cells were compromised for *de novo* folding, post-stress protein refolding and in regulated degradation of a model terminally misfolded protein. Together these findings pinpoint Hsp70 as a key link between oxidative stress and proteostasis, information critical to understanding cytoprotective systems that prevent and manage cellular insults underlying complex disease states.

## Introduction

Protein molecular chaperones facilitate proper conformational folding of nascent polypeptides, refolding of proteins from a misfolded state, and shuttling of protein substrates marked for degradation to proteolytic machinery. Disruption of proteomic management can result in accumulation of non-functional and/or aggregated proteins, which are often implicated in adverse neurological conditions such as Alzheimer’s disease, Parkinson’s disease, and Huntington’s disease ^1–3^. In both humans and the yeast *Saccharomyces cerevisiae*, a key chaperone group modulating many of these tasks are the cytosolic Hsp70 chaperone proteins, composed of an amino-terminal nucleotide binding/ATPase domain (NBD) and a carboxyl-terminal substrate binding domain (SBD) that communicate through allosteric interactions with assistance from other highly conserved chaperones (Hsp40, which assists in Hsp70/substrate contact; and Hsp110, which assists in nucleotide exchange) in a process termed chaperone cycling ^4,5^. These co-chaperones serve to regulate the rate of intrinsic nucleotide hydrolysis and substrate binding and release of Hsp70. Binding of a substrate within the SBD, along with stimulation by Hsp40, induces a conformational cascade that accelerates ATP hydrolysis within the NBD ^6^. ATP binding, hydrolysis and release (nucleotide cycling) activity thus alters affinity of the substrate binding domain for its targets and also modulates the iterative Hsp70 release/binding mechanism that promotes protein folding. *S. cerevisiae* expresses four cytosolic Hsp70s, of which Ssa1 (human homolog HSPA1A) is the most abundant, constitutively expressed homolog. Yeast additionally possesses another constitutively expressed homolog (Ssa2), and two stress-induced homologs (Ssa3/4), whose expression is restricted until exposure to environmental stress ^7^. However, expression of any one of the four Ssa isoforms is sufficient to allow viability ^8,9^.

Reactive oxygen species (ROS) are unstable and highly reactive molecules derived from oxygen, known to be particularly detrimental to lipids, DNA, and proteins ^10^. ROS are a common byproduct of normal cellular functions and are typically present at low concentrations, but uncontrolled/acute increases in abundance can negatively impact cell physiology ^11^. Uncontrolled ROS has been implicated as a contributing factor in aging, cancer, and neurodegenerative disease ^12,13^. Oxidative stress management is particularly important for protein homeostasis, as evidenced by the vulnerability of proteins at all stages of their life cycle, and especially nascent polypeptides ^14^. Several chaperones have been found to act as sensors for oxidative stress, triggering downstream events that protect the cell from damage to proteins ^15^. Due to their variable reactivity, reversibility, and breadth of oxidative states, cysteines represent a versatile and useful sensor for cellular redox state ^16,17^. Cysteine oxidation of the yeast peroxidredoxin Tsa1 was shown to alter its normal behavior, whereby it switches from a hydrogen peroxide scavenger to a passive “holdase” chaperone, actively recruiting Hsp70/Hsp104 to assist with clearance of misfolded proteins ^16^. In bacteria, Hsp33 has been shown to contain reactive cysteines that sense oxidative stress, activating substrate binding capacity for prolonged periods of time to similarly convert the inactive chaperone into a holdase to prevent substrate aggregation ^18^. Reactive cysteines also play a key role in the regulatory relationship between Gpx3 and Yap1, the major transcription factor of the oxidative stress response in yeast ^19^. In mammalian cells, oxidative stress impacts the regulation of the transcription factor Nrf2 by KEAP, a cysteine-rich protein whose oxidation and subsequent degradation enable Nrf2 to induce an oxidative stress response circuit ^20,21^.

Cysteines within Hsp70s have been demonstrated to be important for stress defense, spanning various organisms, cellular compartments, and thiol-modifying compounds ^22–24^. The triterpenoid compound celastrol, used in traditional Chinese medicine to reduce inflammation, is a thiol-modifying compound that was shown to activate the major regulator of the heat shock response (HSR) in yeast, Hsf1 ^25^. The heat shock response is a general stress response, activating the transcription of hundreds of downstream genes, due to several types of stress ^26^. Ssa1 contains two reactive cysteines (C264 and C303), located within the nucleotide binding domain, and nullification of these cysteines via serine substitution was shown to abolish Hsf1 reactivity to thiol-modifying compounds and an inability to activate the HSR in response to oxidative stress, but not thermal stress ^27^. Conversely, mimicking cysteine oxidation to a sulfinic acid moiety via substitution of both cysteines with aspartic acid resulted in a constitutively hyperactive HSR ^27^. Recently, we and others characterized the mechanistic repression of Hsf1 by Ssa1 through direct physical interaction, establishing a correlation between Hsp70 functional status and induction of the heat shock response as an effort to relieve proteotoxic stress ^28,29^. However, a definitive relationship between thiol modification of Ssa1, its regulation of the HSR through Hsf1, and downstream effects on general proteostasis remains to be determined.

Modelling thiol oxidation through genetic techniques has allowed several groups to probe the effects of cysteine modification on chaperone proteins without the broad off-target effects of introducing exogenous oxidative compounds. Distinct amino acid substitutions, such as aspartic acid, were used to effectively mimic the steric and electrostatic changes that result from thiol oxidation. Oxidomimetic substitutions in the human inducible Hsp70 (HSPA1A) resulted in structural changes to the nucleotide binding pocket of the chaperone, resulting in both enzymatic and functional deficiencies ^30^. Oxidomimetic mutations also modified the behavior of the endoplasmic reticulum-localized Hsp70 protein BiP from that of a protein-maturation assisting ‘foldase’ into a less enzymatically active holdase state, where substrates are passively bound without regulated release. This chaperone reprogramming increased cell survival under conditions of oxidative stress, but negatively impacted growth under non-stress conditions ^31^. While increased association of this mutant form of BiP (C63D) with unfolded polypeptide substrates offered a protective effect against oxidative stress, a C63H substitution that mimics the bulkiness but not the charge of a sulfinic acid was shown to have a dominant negative effect in non-stress conditions, likely due to increased holdase activity that prevented proper folding and secretion of proteins ^32^. The enzymatic effects of mimicked oxidation through amino acid substitution mirror the reductions in nucleotide binding, ATPase activity, and protein refolding seen from exogenously added oxidizing or alkylating compounds ^30,33,34^. In HeLa cells, glutathionylation of Hsp70 cysteines caused aberrations in substrate recognition and binding, again supporting the notion that Hsp70 cysteines are both highly reactive and functionally vulnerable residues ^24^.

To better understand the roles of C264 and C303 within Ssa1 as oxidative stress sensors and targets, as well as downstream implications of thiol stress on general Ssa1-dependent proteostasis, we undertook a combined biochemical and genetic approach. The dual sulfinic acid mimic Ssa1-2CD (C264D, C303D) was impaired for several critical Hsp70 functions *in vitro*, including nucleotide binding, ATP hydrolysis, and the refolding of chemically denatured protein. Exogenous oxidation of Ssa1 with hydrogen peroxide mirrored the deficiencies of the Ssa1-2CD mutant, but the cysteine-null Ssa1-2CS (C264S, C303S) variant was impervious to exogenous oxidation, confirming cysteines 264 and 303 as the primary relevant sites of oxidative modification. The *ssa1-2CD* mutant was found to exert a dominant negative growth phenotype in cells and was additionally unable to facilitate growth as the sole cytosolic *SSA* isoform. The oxidomimetic *ssa1-2CD* mutant also exhibited reduced ability to maintain general proteostasis *in vivo*, as evidenced by defective folding and refolding of the reporter protein FFL-GFP as well as inability to promote degradation of a chronically misfolded protein. Extending previous work, we demonstrate that Ssa1-2CD failed to associate with Hsf1 and repress its activity under non-stress conditions, resulting in chronic activation of the HSR. Taken together, these results support a model wherein cysteines within the primary, constitutive cytosolic Hsp70 chaperone are subject to oxidative modification that negatively impacts general proteostasis but also concomitantly engages the HSR to promote a cytoprotective response.

## Results

### Ssa1 oxidation reduces ATP binding and hydrolysis

The conformational changes between the NBD and SBD that allow Hsp70 chaperones to iteratively bind and release substrate are dependent on allosteric signals from interactions with nucleotide, substrate, and co-chaperones. This process was previously shown to be disrupted by exogenous treatment with thiol-reactive compounds ^33^. The sulfhydryl alkylating reagent *N*-ethylmaleimide negated the ability of yeast Ssa1 to bind ATP-agarose, as well as to hydrolyze ATP ^34^. To continue exploring the Hsp70/nucleotide relationship and to confirm the relevant amino acid targets of oxidative attack, we generated and purified to homogeneity recombinant Ssa1 proteins with the wild type sequence, serine substitutions (Ssa1-2CS), or aspartic acid substitutions (Ssa1-2CD) at cysteines 264 and 303 ^27,35^ (Fig. S1A). Because binding of ATP within the NBD generates an allosteric signal to induce conformational change of Hsp70, we hypothesized that modification of C264 and C303 would alter the ability of Ssa1 to interact with nucleotide. To test this, we first measured nucleotide binding ability using ATP-agarose chromatography. Purified proteins were incubated with ATP-agarose, followed by several washes and elution with sample buffer and immunoblot. Comparable amounts of Ssa1 (WT) and Ssa1-2CS were eluted from the ATP-agarose beads after incubation, while the Ssa1-2CD protein was unable to bind to the same extent (Fig. 1A, quantitated in Fig. 1B). Exogenous addition of 1 mM hydrogen peroxide prior to incubation with the beads significantly reduced the signal of eluted Ssa1, while Ssa1-2CD ATP binding remained low and nearly identical to untreated sample (Fig. 1B). Importantly, Ssa1-2CS retained full ATP binding capacity regardless of hydrogen peroxide treatment, suggesting that reduced nucleotide binding is a functional consequence of cysteine oxidation.

**Fig. 1.**
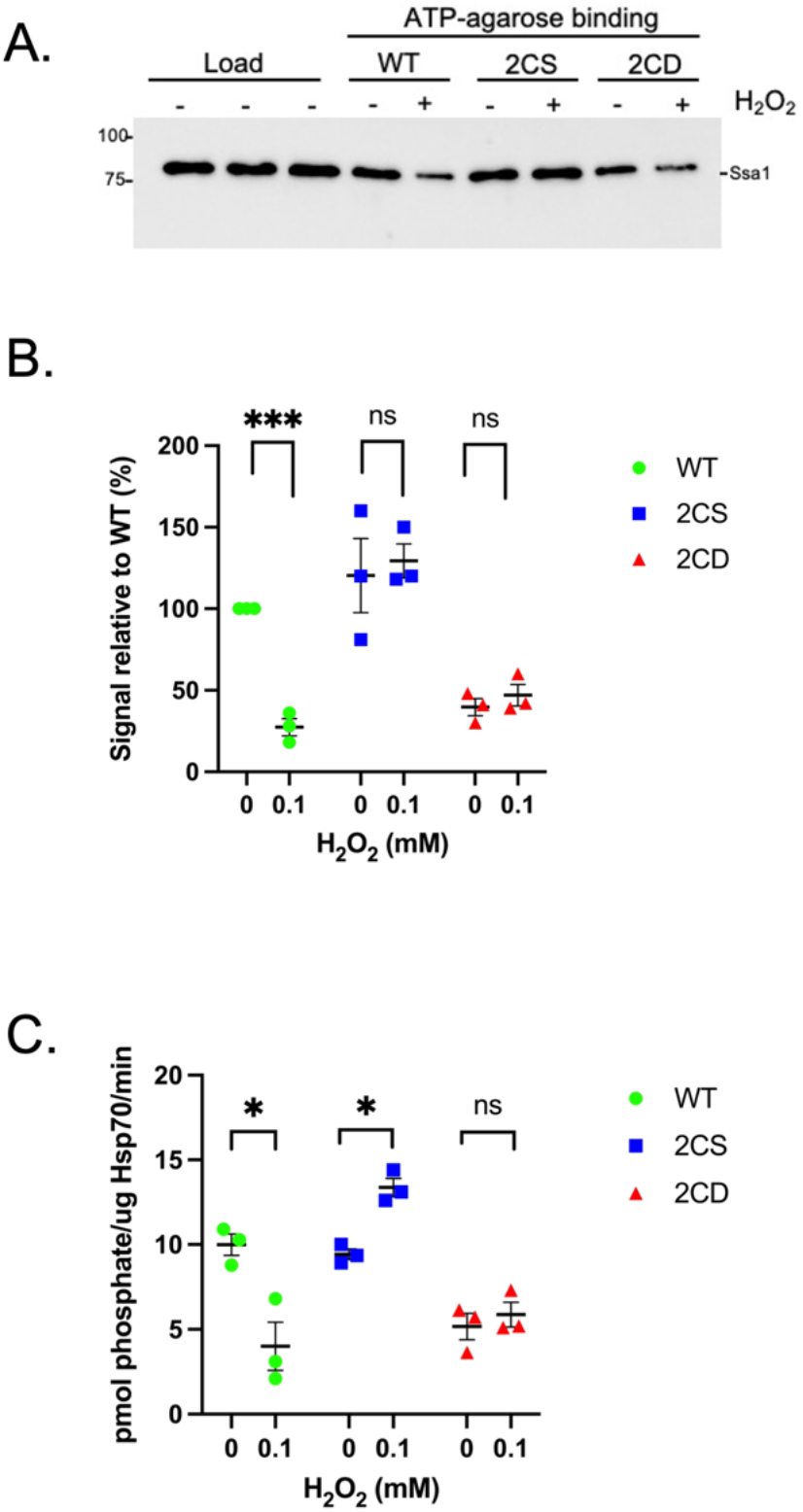
Nucleotide binding and hydrolysis are similarly impaired by oxidomimetic substitution in Ssa1-2CD and exogenous oxidation of Ssa1. (A) Immunoblot displaying elution fraction of 25 uL ATP-agarose bead volume incubated with 1 uM total respective protein treated or not with 1 mM hydrogen peroxide. (B) Quantification of the signal in *A* converted to relative percentage of untreated Ssa1 and normalized for load. (C) Rate of ATP hydrolysis by 0.1 uM of respective proteins, treated or not with 1 mM hydrogen peroxide. Bolded horizontal bars indicate mean, and error bars indicate SEM.

To assess whether the reduced binding interaction resulted in a downstream change in catalytic activity, we tested the ability of each isolated protein to hydrolyze ATP using a malachite green assay to quantify released phosphate. Akin to nucleotide binding, Ssa1 and Ssa1-2CS exhibited comparable levels of specific activity, while Ssa1-2CD demonstrated significantly reduced hydrolysis. (Fig. 1C). Treatment with hydrogen peroxide also drastically reduced ATP hydrolysis of the Ssa1 but not Ssa1-2CS protein, while Ssa1-2CS retained full and even slightly elevated activity. Hydrolysis of ATP in the NBD is stimulated by Hsp40. We therefore wanted to determine if stimulation by Hsp40 could overcome the basal hydrolysis deficit of Ssa1-2CD. Interestingly, the addition of the Hsp40 Ydj1 equally stimulated all Ssa1 isolates, but was not able to rectify the deficiency in hydrolysis from Ssa1-2CD (Fig. S1B). Together, these data indicate that modification of cysteines, by both exogenous oxidation and oxidomimetic mutation, results in an altered relationship between Ssa1 and nucleotide resulting in an inability to bind and hydrolyze ATP.

### Ssa1 oxidation inactivates substrate folding

Disruption of nucleotide interaction within the NBD of Hsp70 has negative implications for the allosteric conformational changes signaled through the linker to the SBD ^36^. We hypothesized that this disruption in signal would affect the Hsp70/substrate relationship. To determine the consequences of impaired nucleotide interaction on protein folding by Hsp70, we utilized recombinant firefly luciferase (FFL) as a substrate for Ssa1. Susceptible to chemical denaturation, the enzymatic activity of properly folded FFL to produce chemiluminescence in the presence of the substrate luciferin has been previously used to measure substrate refolding by yeast chaperones *in vitro* ^37–39^. We therefore reconstituted the Hsp70 folding triad that includes Hsp70 (Ssa1), Hsp40 (Ydj1), and Hsp110 (Sse1) to examine refolding of chemically denatured FFL *in vitro*. Ssa1 and the cysteine-null Ssa1-2CS were found to have comparable basal ability to refold FFL post-exposure to guanidinium hydrochloride, while the yield of active enzyme produced by Ssa1-2CD was dramatically reduced (Fig. 2A). The yeast disaggregase Hsp104 assists Hsp70 and Hsp40 in the reactivation of aggregated proteins ^40^. After addition of this disaggregase to our chaperone mixture, we found that while Hsp104 approximately doubled the final yield of FFL recovered for all strains, there was still a significant defect in folding by Ssa1-2CD as compared to Ssa1-WT and Ssa1-2CS (Fig. 2B). To determine the effects of exogenous oxidation on refolding, we measured the final yield of recovered FFL after pre-exposure of Ssa1 alone to 1 mM hydrogen peroxide, and found that there was a significant decrease in recovery by SSA1-WT after oxidation, while Ssa1-2CS and Ssa1-2CD interestingly exhibited increased folding with respect to their untreated matched samples (Fig. 2C). We hypothesize that this may be due to a secondary effect that shifts the conformation of Ssa1 in a way that is favorable for folding. However, because the observed increases occurred in both the serine and aspartic acid mutants, these effects are likely independent of cysteines 264 and 303.

**Fig. 2.**
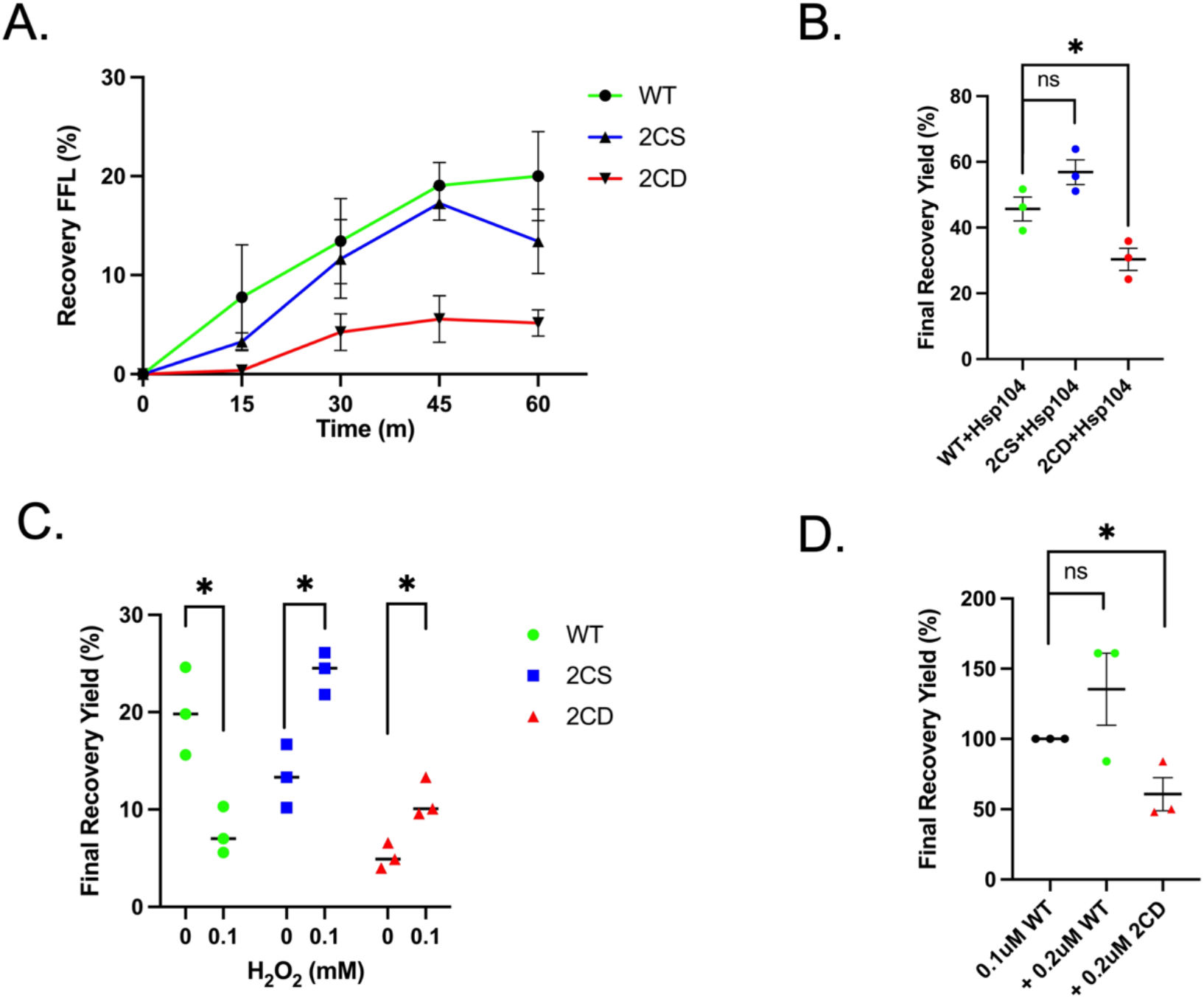
Mimicked and exogenous thiol oxidation negatively impact *in vitro* protein refolding. (A) Firefly luciferase (FFL) recovery over time in the presence of 0.1 uM of the indicated Ssa1 protein and co-chaperones Ydj1 (0.2 uM) and Sse1 (0.05 uM), as described in detail in *Materials and Methods*. (B) End point measurements of folding reactions including addition of 1 uM Hsp104. (C) End point measurements of folding reactions using Ssa1 variants treated or not with 1 mM hydrogen peroxide. (D) End point measurements of folding reactions, indicating the respective protein additionally added to reactions as in (A). Bolded horizontal bars indicate mean, and error bars indicate SEM.

In at least two prior reports, thiol modification or oxidomimetic substitution in Hsp70 chaperones has generated increased holdase capacity, binding substrate without ATP-regulated release, and therefore resulting in a dominant negative effect on protein refolding ^32,33^. To assess the possibility that a C264/C303 oxidomimetic displays similar dominant negative properties, we titrated 0.2 uM additional Ssa1 or Ssa1-2CD into a pre-existing chaperone cocktail containing 0.1 uM Ssa1. We found that compared to Ssa1 alone, the Ssa1/Ssa1-2CD pool was hindered in refolding ability (Fig. 2D). This suggests that the Ssa1-2CD oxidomimetic may non-productively bind substrate and inhibit refolding in a dominant negative manner. Altogether, these results substantiate the negative functional effects of thiol oxidation on protein folding by Ssa1.

### ssa-2CD is incapable of functioning as the sole SSA isoform and is dominant negative

To complement our *in vitro* studies, we addressed consequences of Ssa1 oxidation through the genetic cysteine null and oxidomimetic Ssa1 mutants. We initially attempted to express *ssa1-2CD* as the sole cytosolic *SSA* gene in a quadruple *ssa1Δssa2Δssa3Δssa4Δ* deletion background ^41^. This strain contained a wild type copy of *SSA1* on a *URA3*-selectable plasmid and was additionally transformed with a *HIS3-*selectable plasmid expressing wild type *SSA1* or the *ssa1-2CS* or *ssa1-2CD* mutants. A plasmid shuffle technique was used to selectively isolate colonies that possessed only the *HIS3* plasmid via plating on 5-fluoroorotic acid (5FOA) media. Surprisingly, all recovered colonies expressing *SSA* alleles grew at identical rates when plated (Fig. S2A), inconsistent with the slow-growth phenotype previously reported for the *ssa1-2CD* mutant ^27^. Sequencing of the recovered *HIS3*-marked plasmid revealed that the *ssa1-2CD* allele had converted to the wild type sequence encoding the original cysteine residues (Fig. S2B). We hypothesize that this was due to a recombination event between the two plasmids, whereby the likely inability of *ssa1-2CD* to function as the sole expressing cytosolic *SSA* gene resulted in selection for rare allele exchange events (Fig. S2C). After several attempts resulting in either no viable colonies or only allele-exchanged colonies, we concluded that the *ssa1-2CD* allele is incapable of sustaining viability as the sole cytosolic *SSA* isoform due to the functional defects demonstrated in Figs. 1 and 2.

We elected to continue with oxidomimetic expression *in vivo* in an *ssa1Δssa2Δ* deletion background, where Ssa3/4 are present at low levels to support viability, but cell growth is still significantly impaired in non-stressed conditions ^27^. We additionally wanted to ensure that expression of the wild type and mutant alleles best represented natural levels and examined different heterologous promoters for suitability. We found that *SSA1* allele expression from the low-expressing *CYC1* promoter on centromeric *(CEN)* yeast expression vectors resulted in diminished growth for all genotypes, suggesting general Ssa protein insufficiency. However, expression from the stronger CEN-*TEF* vector backbone resulted in normal growth for *SSA1* and *ssa1-2CS* strains, while the *ssa1-2CD* and empty-vector control both grew at rates similar to each other and consistent with previous reports (Fig. 3A, B and Fig. S3A) ^27^. Immunoblots confirmed that Ssa1 protein levels were expectedly lower driven from the *CYC1* promoter in *ssa1Δ ssa2Δ* cells as compared to the DS10 parent strain (Fig. S3B,C). Intriguingly, *TEF-*driven Ssa1 protein levels were similar between *SSA1, ssa1-2CS*, and the parent DS10 strains, but the *ssa1-2CD* allele was clearly produced lower levels of Ssa1-2CD protein (Fig. S3B,C). We reasoned that Ssa1-2CD expression was either being actively curtailed or that the mutation resulted in a protein more susceptible to degradation. To test protein stability, we treated cells with the protein translation inhibitor cycloheximide and tracked the existing pool of Ssa1 from each allele by immunoblot, finding that Ssa1-2CD was stable over the course of 3 hr (Fig. S3D). These results led us to the conclusions that steady state Ssa1-2CD levels might be restricted in actively growing cells but that the protein itself was not inherently destabilized relative to wild type Ssa1.

**Fig. 3.**
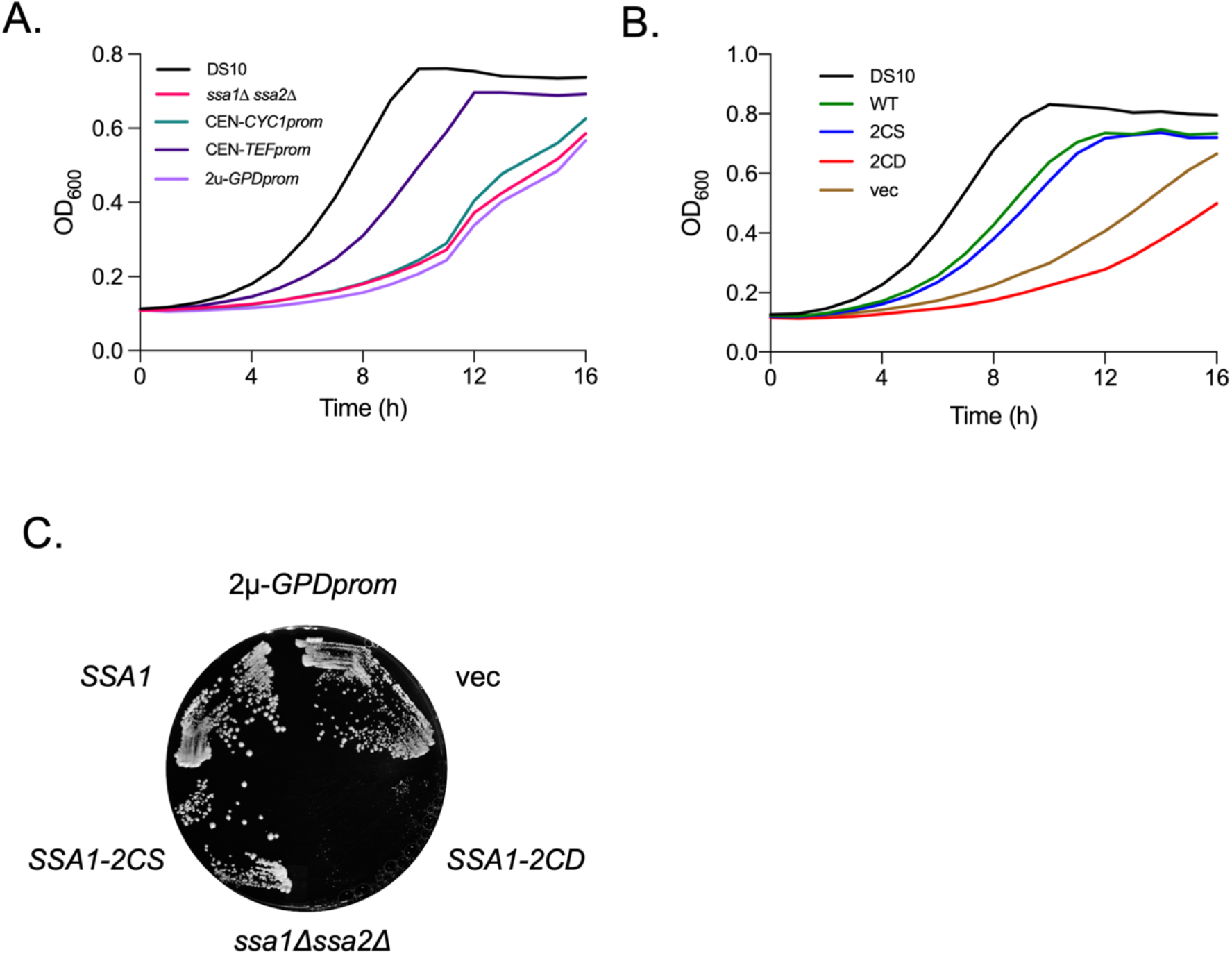
The thiol oxidomimetic *ssa1-2CD* allele displays dominant negative growth impairment. (A) 16-hour growth curve of parent strain DS10, *ssa1Δ ssa2Δ*, and wild type *SSA1* driven by the indicated promoter in a *ssa1Δ ssa2Δ* background. (B) 16-hour growth curve of DS10 and indicated *SSA1* allele expression driven by the *TEF* promoter, in an *ssa1Δ ssa2Δ* background. (C) 48-hour plate growth of each indicated *SSA1* allele expressed from the 2µ pRS423TEF vector backbone, in an *ssa1Δssa2Δ* background.

Our in vitro experiments indicated that *ssa1-2CD* exerted dominant negative effects on protein folding. We noted that culture growth rates of the *ssa11-2CD* strain were slower than even the empty vector *ssa1Δ ssa2Δ* control strain, suggesting that expression of the Ssa1-2CD protein at even moderate levels was more detrimental than having no Ssa1 at all (Fig. 3B). To further explore this phenomenon, we expressed all *SSA1* alleles from the strong *GPD* promoter on a 2µ vector backbone ^42^. This level of overexpression of *SSA1* and *ssa1-2CS* reduced growth rates to a level similar to the empty vector background control, consistent with previous reports that chaperone overexpression can be deleterious (Fig. 3C) ^43^. However, overexpression of the *ssa1-2CD* allele resulted in near-total cessation of growth despite the presence of Ssa3/4 in this background, indicating overexpression toxicity beyond the inability to complement loss of Ssa1/2 functions.

### Ssa1-2CD fails to physically associate with Hsf1 to repress the HSR

Ssa1 has been recently shown by our laboratory and others to act as a repressor of the heat shock response transcription factor Hsf1 through physical association with both the amino- and carboxyl-terminal transcriptional activation domains ^28,29,44^. We have also previously published that thiol-modifying compounds induce a heat shock response in *SSA1*, but not *ssa1-2CS*, cells ^27^. Consolidating these findings, we hypothesized that thiol stress alters C264 and C303 within Ssa1, inactivating the chaperone resulting in release of Hsf1 and subsequent induction of the HSR. We first confirmed that the HSR was chronically activated in our *ssa1-2CD* strain utilizing an Hsf1-responsive HSE-lacZ reporter (Fig. 4A). *ssa1-2CS* cells exhibited appropriate HSR repression, verifying that the endogenous cysteines are not required for Ssa1 to function as a repressor of Hsf1. To test our hypothesis that Ssa1-2CD is defective in Hsf1 association, Hsf1-GFP-FLAG was co-expressed in cells containing either *CYC1-*driven *SSA1* and *ssa1-2CS* or *TEF*-driven *ssa1-2CD* alleles to control for differential Ssa1 protein levels, and co-immunoprecipitations were performed. Both Ssa1 and Ssa1-2CS associated with Hsf1, but no detectable signal was observed for Ssa1-2CD (Fig. 4B). None of the Ssa1 proteins were found to associate with the GFP-FLAG control, confirming specificity of Hsf1 binding. These data indicate that the oxidomimetic Ssa1-2CD is unable to productively bind the bipartite contact sites on Hsf1 and provide a molecular mechanism to explain oxidative stress sensing by Hsf1 via Cys264/303 of Ssa1.

**Fig. 4.**
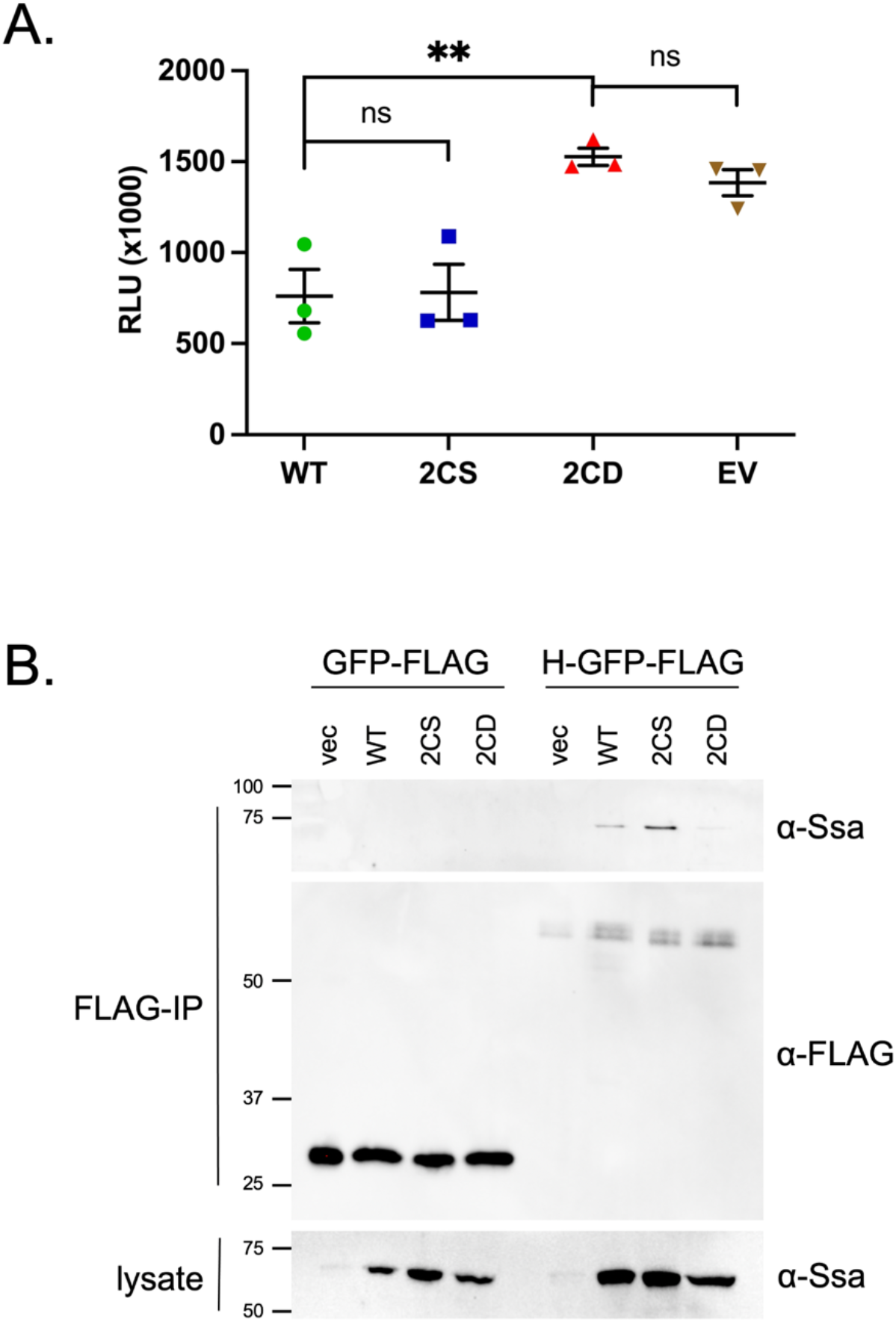
Ssa1-2CD fails to bind and repress the heat shock regulator Hsf1. (A) HSE-lacZ activity of strains containing wild type and mutant *SSA1* alleles grown at 30*°*C. To normalize Ssa1 protein levels, *SSA1* and *ssa1-2CS* were expressed from the CYC1 promoter and *ssa1-2CD* from the stronger TEF promoter. (B) Co-immunoprecipitation of tagged Hsf1-GFP-FLAG or GFP-FLAG control and the indicated Ssa1 proteins. Bolded horizontal bars indicate mean, and error bars indicate SEM.

### ssa1-2CD is defective in protein folding, solubilization and refolding in vivo

Hsp70 proteins are critical factors for ensuring proteostasis ^5^. We hypothesized that mimicked cysteine oxidation would have negative consequences for Hsp70 protein surveillance activities. Data presented in Fig. 2 demonstrate that the Ssa1-2CD protein or exogenously oxidized Ssa1 are defective in protein folding. To complement the *in vitro* folding assays, we co-expressed with the *SSA1* alleles a previously generated and well documented firefly luciferase (FFL)-GFP fusion protein known to require the Hsp70 chaperone system for folding in living cells ^45,46^. To monitor *de novo* folding of nascent polypeptides, we utilized the methionine-repressible promoter of the FFL-GFP plasmid to induce expression of FFL in log phase cells. FFL activity was measured by luciferase assay over the course of 90 min. The *ssa1-2CS* mutant was able to fully complement the *de novo* folding defect observed in *ssa1Δ ssa2Δ* cells relative to cells expressing *SSA1* while the *ssa1-2CD* mutant was significantly defective (∼ 60% of wild type luciferase activity) (Fig. 5A). To account for the lower abundance of Ssa1-2CD, we also examined *de novo* folding activity in a *CYC1-SSA1* strain, and found that this reduced level of expression still maintained higher FFL activity than observed in the *TEF-ssa1-2CD* background, indicating that absolute protein levels do not explain the reduced capacity for FFL folding seen with Ssa1-2CD (Fig. S3E).

**Fig. 5.**
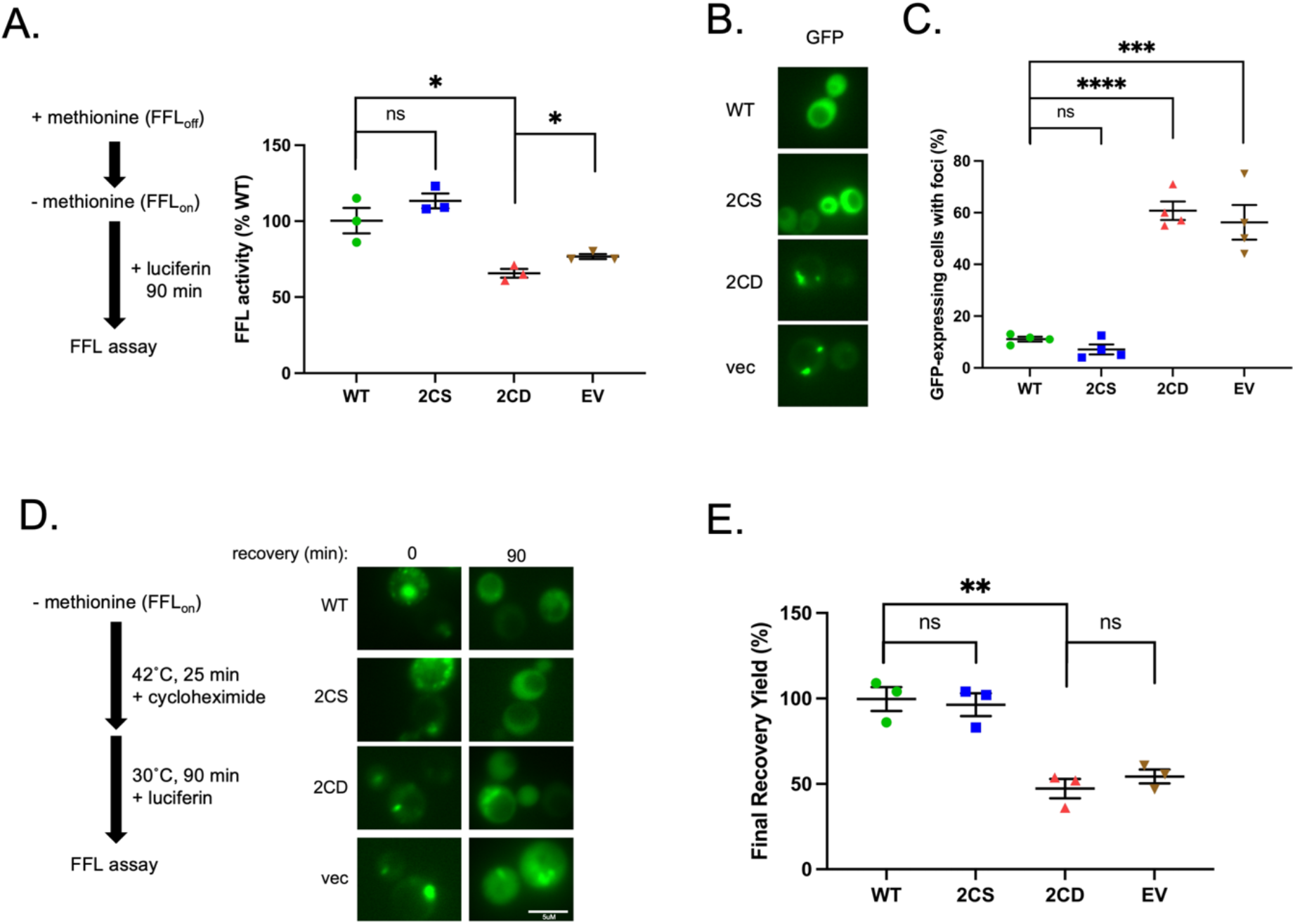
The *ssa1-2CD* mutant exhibits multiple deficiencies in proteostasis. (A) End point *de novo* folding ability of *SSA1* or mutant *ssa1* alleles as measured by luciferase activity assay, monitored over 90 min. (B) Images representing FFL-GFP fluorescence in *SSA1* or *ssa1* mutant strains grown overnight at 30*°*C. (C) Quantification of (B) in terms of foci per cell. Each data point represents percentage of cells containing foci from a minimum of 50 cells per replicate. (D) Schematic of FFL-GFP recovery assay and representative micrographs detailing *SSA1* and mutant *SSA1* strains at 0 and 90 min after initiation of cycloheximide chase. (E) Quantification of assay detailed in (D), measured as the percentage FFL-GFP chemiluminescence activity at t_90_ in *SSA1* and respective ssa1 mutant strains, relative to the same sample at t_0_. Bolded horizontal bars indicate mean, and error bars indicate SEM.

Misfolded proteins are known to aggregate in concert with protein chaperones, including sequestrases and disaggregases, and can be visualized *in vivo* using fluorescent tagging approaches ^47,48^. To examine the status of properly folded and potentially misfolded FFL, overnight cultures of *SSA1, ssa1-2CS, ssa1-2CD* and a vector control containing the FFL-GFP expressing plasmid were visualized using fluorescence microscopy. Micrographs displaying representative images show that FFL-GFP was found to be fully soluble in *SSA1* and *ssa1-2CS* strains while *ssa1-2CD* and the vector control exhibited large foci, visible as fluorescent puncta that were present in a significantly higher percentage of cells (Fig. 5B,C). As an additional orthogonal approach to complement our *in vitro* findings, we determined the ability of each strain to refold heat-denatured substrate. Log-phase cells were washed to remove methionine and induce FFL-GFP expression to generate a pool of substrate. Cycloheximide was then added to prevent additional FFL-GFP expression and steady-state luminescence activity was measured. Cellular FFL-GFP was denatured by incubating cultures at 42°C, followed by a recovery period at 30°C. Cells were then visualized by fluorescence microscopy and luminescence was determined. All strains contained puncta immediately after heat shock, and while *SSA1* and *ssa1-2CS* strains resolved FFL-GFP aggregates, *ssa1-2CD* and the empty vector control failed to do so (Fig. 5D). These results were mirrored when FFL enzymatic activity was examined, with the *ssa1-2CD* and the empty vector control only managing to restore approximately 50% of original pre-heat shock FFL-GFP activity (Fig. 5E). Taken together, these data indicate that the mimicking of chronic oxidation of Ssa1 cysteines 264 and 303 dramatically undermines general proteostasis, with negative consequences for the proper folding of nascent translating polypeptides and the re-folding of stress-denatured proteins.

### ssa1-2CD cells fail to clear permanently misfolded proteins from aggregates

Chaperones, most notably the Hsp70 system, are tightly integrated into the protein quality control system and selectively regulate protein degradation via presentation of substrate to ubiquitin ligases ^49^. To investigate how thiol modification of Ssa1 cysteines affects degradation of misfolded proteins, we utilized a well-characterized substrate, tGnd-GFP ^49,50^. This artificial construct contains a truncated version of the Gnd1 protein fused to GFP that results in an unfoldable substrate with defined kinetics of degradation through the ubiquitin-proteasome pathway. We first expressed tGnd-GFP in the presence of the *SSA1* alleles and observed a greater than two-fold increase in steady state levels of the substrate by immunoblot in the *ssa1-2CD* and vector control strains (Fig. 6A, 6B). To determine the status of accumulated tGnd-GFP, log phase cells were treated with cycloheximide to stop further synthesis and imaged by fluorescence microscopy. Representative micrographs show the presence of tGnd1-GFP foci presence in all strains at t^0^, indicating that misfolded protein was sequestered into protein aggregates (Fig. 6C). After 90 min, foci remained in a significantly higher amount of *ssa1-2CD* and vector control cells compared to *SSA1* and *ssa1-2CS* cells, where foci were largely eliminated, indicative of impaired substrate processing and degradation (Fig. 6C,D). Taken together, these data support the conclusion that Ssa1 cysteine oxidation, as mimicked by aspartate substitution, renders the chaperone defective in promoting the degradation of terminally misfolded substrates.

**Fig. 6.**
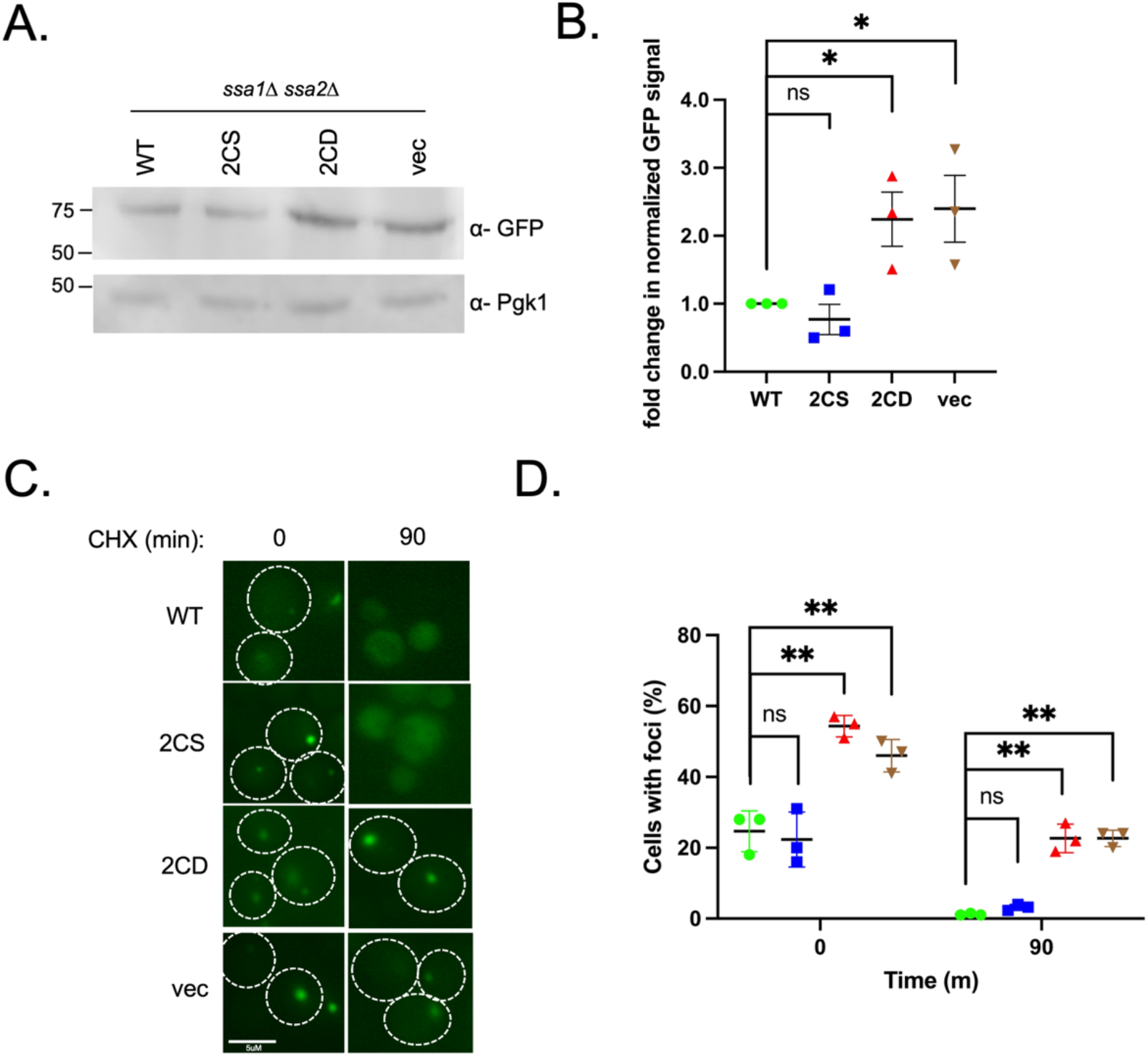
Degradation of the misfolded protein tGND-GFP is chronically impaired in *ssa1-2CD* cells. (A) Western blot analysis of steady state levels of the chronically misfolded protein tGND, co-expressed with *SSA1* or mutant *ssa1* alleles taken from cells in log phase. (B) Quantification of signal in A, measured as fold change of each mutant allele relative to *SSA1*, from three replicate blots. (C) Representative images of cycloheximide chase, monitoring tGND-GFP foci presence over 90 min for each respective allele. (D) Quantification of images in C, with each data point representing a percentage of cells containing foci from a minimum of 50 cells per replicate. Bolded horizontal bars indicate mean, and error bars indicate SEM.

## Discussion

In this study, we explored in detail the functional consequences of oxidation of two key cysteines in Ssa1 previously demonstrated to be required for activation of the HSR by thiol-reactive molecules via genetic and chemical approaches. Analysis using purified proteins revealed that Ssa1-2CD, but not the Ssa1-2CS mutant, is compromised for ATP binding and hydrolysis, as well as ATP-dependent protein folding. Nearly identical results were observed upon treatment of wild type Ssa1 protein with hydrogen peroxide, validating the utility of the aspartic acid substitution to mimic the sulfinic acid state of the cysteine thiol group after oxidation. Importantly, the non-reactive cysteine null substitution to serine (Ssa1-2CS) was impervious to hydrogen peroxide at concentrations that impaired Ssa1 enzymatic activity *in vitro*. We can therefore conclude that cysteines 264 and 303 are exclusively responsible for the observed behaviors in vitro and in vivo, at least under our experimental conditions. In addition to these two cysteines in Ssa1 located on lobe IIB within the nucleotide binding domain, a third cysteine residue is conserved in most Hsp70 homologs deeper within the nucleotide binding cleft (C15 in Ssa1). Our results suggest that C15 is not a critical target for oxidation for Ssa1. This finding contrasts with previous work from the Sevier laboratory where C63 of the endoplasmic reticulum-resident Hsp70 Kar2/BiP was found to be sensitive to oxidation ^51^. However, both yeast Kar2 and mammalian BiP lack homologous cysteines 264 and 303, complicating a direct comparison. Notably, all three cysteines are sensitive to modification by the strong alkylating agent N-ethylmaleimide (NEM); NEM-Ssa1 also exhibited the same range of functional defects as our oxidomimetic mutant and peroxide-treated Ssa1 ^33^. NEM-treated Ssa1 displayed altered trypsinization profiles and intrinsic tryptophan fluorescence indicative of structural deformation within the NBD ^34^. In human Hsc70, it was found that modification of nucleotide pocket-facing C17 displaced a catalytic magnesium ion that is crucial for nucleotide hydrolysis by increasing the distance of that magnesium from several interacting residues ^52^. These results are also consistent with previous data that methylene blue oxidizes cysteines 267 and 306 in the stress-inducible human Hsp70, resulting in reduced ATP binding and hydrolysis ^30^. *in silico* modeling in this report suggested that oxidation occurs step-wise, where C306, located on the outer surface of lobe IIB of the NBD, is oxidized first, resulting in a conformational change that displaces the inner lobe cysteine, C267, into a solvent-exposed cleft. This structural deformation may increase the accessibility or reactivity of the thiolate anion of cysteine 267 and ultimately disfigure the entire NBD to reduce nucleotide binding affinity and dependent enzymatic activities. This model is in complete agreement with a previous study of ours demonstrating that reaction of C264 in Ssa1 with a 4-hydroxynonenal alkyne and subsequent derivatization via click chemistry was lost in a C303S mutant ^27^.

We found that addition of Ssa1-2CD inhibited productive firefly luciferase refolding by Ssa1. This result is consistent with previous data that NEM-Ssa1 likewise blocked Ssa1-dependent folding but not the ability to inhibit aggregation of misfolded protein. Indeed, oxidation, alkylation, or substitution with a bulky side chain residue of C63 in Kar2 significantly enhances the “holdase” ability of that Hsp70 ^51^. These findings are all consistent with a model wherein oxidation disrupts productive allosteric communication between the NBD and SBD but does not affect, and may even enhance, polypeptide binding by the SBD, resulting in competition for Hsp70 binding sites and restricting productive folding reactions. Such a model would explain the semi-dominant negative growth phenotypes we observed upon expression of *ssa1-2CD* in cells that still retained expression of the inducible *SSA* isoforms Ssa3 and Ssa4. Importantly, this model also provides a mechanism by which oxidation of only a portion of the highly abundant pool of Ssa1 and Ssa2 could exert deleterious effects on cell functions and growth, as it is highly unlikely that transient exposure to oxidants would modify the majority of available Hsp70. Unlike the findings of Wang and coworkers, wherein oxidized Kar2 renders cells hyper-resistant to oxidative stress, we found no evidence for any gain of function phenotypes in the *ssa1-2CD* strain ^51^.

As demonstrated by substitution with serine or alanine residues, cysteines 264 and 303 in Ssa1 are dispensable for growth at elevated temperatures, as is C63 in Kar2/BiP ^51^. However, the cysteines are essential for activation of stress responses and cytoprotection by cells exposed to oxidizing compounds ^27,51^. This pattern is indicative of a residue that has been evolutionarily conserved to sense and react to an oxidative stressor. The reversibility of cysteine modification is especially useful as a signal, as it allows for chaperones that contain them to switch between conformations that may have different purposes, with the additional involvement of redox management systems such as the thioredoxin and glutathione pathways ^53^. A key example of the utility of Ssa1 cysteine signaling is the transient activation of the HSR upon treatment with alkylating or oxidizing agents. Because yeast Hsf1 lacks cysteines, it is unable to directly respond to such insults in contrast to human HSF1 that is directly modified on exposed cysteines to activate the HSR to rebalance proteostasis ^54^. C264/C303 of Ssa1 therefore play an elegantly analogous role of oxidative sensor in yeast via the recently described direct repression of Hsf1 by the chaperone. We now for the first time connect these two concepts by showing that Ssa2-2CD fails to bind either known site on Hsf1 in vivo, resulting in a chronically elevated HSR. Interestingly, Ssa3/4 do not contain the cysteine found in the outer face of lobe IIB, C303, only the pocket-facing cysteine C264. Following the *in silico* model, this makes Ssa3/4 less vulnerable to oxidation, which we believe is advantageous in a stressed environment. Ssa3/4 are expressed at much lower levels relative to Ssa1/2, but are upregulated under stress exposure ^7^. One can therefore envision a scenario wherein upon oxidative stress, Ssa1 is inactivated, Hsf1 is liberated and a potent HSR ensues resulting in upregulation of Ssa3/4. These inducible chaperones can then bind to the same sites on Hsf1 to attenuate the HSR and have the added benefit of resisting further oxidative damage/signaling.

We found that multiple aspects of Ssa1 function are compromised in the *ssa1-2CD* strain. Because it would be impossible to deconvolute the pleiotropic effects of whole-cell treatment with exogenous oxidants, a genetic approach using the oxidomimetic substitution is the best approach to understanding functional consequences of Hsp70 modification. We uncovered profound deficiencies in de novo protein folding, extraction of misfolded proteins from aggregates and clearance of terminally misfolded proteins – all salient aspects of proteostasis. Together these phenotypes likely account for the severe slow growth phenotype of the *ssa1-2CD* mutant, its inability to serve as the sole *SSA* isoform and the observed high rate of gene reversion when we attempted the plasmid shuffle approach. Our findings are consistent with and significantly extend previous findings that an oxidomimetic allele of Hsp72 failed to support tau stabilization in human HeLaC3 cells ^30^. Notably, induction of Ssa3/4 via derepression of the HSR is insufficient to restore proteostasis in the *ssa1-2CD* strain but is required to allow viability. It is therefore possible that the *ssa1-2CD* mutant may be entirely dysfunctional in supporting folding and regulated degradation of the proteome but that a reduced level of proteostasis is maintained by Ssa3/4.

Further work is required to fully understand the consequences of oxidative stress on cell and tissue proteostasis as it relates to human health. Many industrial pollutants are oxidants or thiol-reactive molecules (e.g., heavy metals, acroleins) ^55^. Cigarette smoke contains a wide range of thiol-reactive compounds that are known to deplete cellular glutathione levels ^56^ Accumulation of oxidative damage is widely considered to be a key driver of aging, potentially linked to endogenous ROS generated via mitochondrial dysfunction. Indeed, oxidation of a methionine residue in the mitochondrial Mge1 protein, a cofactor of mitochondrial matrix Hsp70, inhibits protein translocation into the organelle resulting in oxidant hypersensitivity ^57^. Clearly more attention must be paid to the impacts of oxidative stress on the protein quality control network going forward.

## Materials and Methods

### Strains, plasmids and yeast cultivation

Yeast strains were derived from either DS10 *(MATa ura3-52 lys1 lys2 trp1-1 his3-11,15 leu2-3112)* or BY4741 (*MATa, his3Δ1; leu2Δ0; met15Δ0; ura3Δ0*) parent strains. The *ssa1Δssa2Δ* strain (*SL314, MATa ura3-52 lys1 lys2 trp1-1 his3-11,15 leu2-3112 ssa1::HIS3, ssa2::LEU2*) was generously provided by the Craig laboratory and is isogenic with DS10 ^8^. Complementation of the lethal *ssa1Δssa2Δssa3Δssa4Δ* strain was conducted using a standard yeast plasmid shuffle technique, with a *URA3*-based *SSA1*-expressing plasmid (a kind gift from the Truman laboratory). The *SSA1* allele plasmids (p413TEF, p413CYC and p423GPD) were constructed by PCR mutagenesis and amplification of the SSA1 ORF using standard cloning methodology with 5’ SpeI and 3’ XhoI restriction sites. All mutants were confirmed using DNA sequencing. All plasmids were transformed into yeast using the rapid yeast transformation protocol ^58^. The FLAG-tagged Hsf1-expressing plasmid was used as previously published ^28^. The pTHD3HA-tGND-GFP plasmid was a kind gift from Dr. Randolph Hampton, University of California, San Diego. The HSE-lacZ plasmid was reported previously ^59^. The p425MET25-FFL-GFP-leu2::URA3 plasmid was used as previously described ^45^. The 6XHis-Smt3-SSA1 plasmid was kindly provided by Dr. Nadinath Nillegoda (Monash University, Australia). *S. cerevisiae* strains were cultured in yeast extract, peptone, dextrose medium (YPD) or synthetic complete (SC) medium (Sunrise Science, San Diego, CA). For growth curve analysis, cells were grown overnight at 30°C. Cells were then sub-cultured and grown to mid-log phase (OD_600_=0.6-0.8), then diluted to OD_600_=0.1 in fresh media. Growth was monitored using a Synergy MX (BioTek, Winooski, VT) microplate reader for 16 hours with shaking at 30°C. Petri plate growth analysis was for two days at 30°C. Cells containing the HSE-*lacZ* reporter were grown to log phase, and diluted to OD_600_ =0.8. 50uL of Beta-Glo reagent (Promega, Madison, WI) and 100 uL of liquid culture was added to each well in a 96-well microplate and measured for beta galactosidase activity using the Synergy MX (BioTek) microplate reader.

### Protein purification

Proteins were isolated as previously described, with several alterations ^35^. Briefly, *SSA1* coding regions were amplified from p413TEF plasmids and subcloned into the pSUMO vector with a 6XHis-Smt3 tag (gifted from the Nillegoda laboratory) ^35,60^. Plasmids were transformed into BL21(DE3) *E. coli* additionally containing the pRARE plasmid and grown overnight in LB Amp/Kan at 37°C. Subcultures were then grown to log phase (OD_600_=0.6) and expression was induced with 0.5 mM IPTG (MilliporeSigma, St. Louis) for 3 hr at 30°C, then cells were collected by centrifugation, washed, and flash frozen for storage at −80°C. The following day, cells were lysed in 30 mL Buffer K (50 mM HEPES-KOH, pH 7.5, 750 mM KCl, 5 mM MgCl_2_) with DNAse, RNAse, protease inhibitors (PI), and 1 mM PMSF. Suspensions were sonicated on ice to lyse, and cell debris was removed by centrifugation for 10 min at 12,500 RCF at 4°C. Supernatant was removed, and the suspension was again centrifuged. The supernatant was brought up to 30 mL with Buffer K and nutated for 1 hr at 4°C with 1 mL bed volume of Buffer K-equilibrated His-Pur cobalt resin (Thermo Scientific), adding 30 uL fresh PI, 1 mM PMSF, and 50 uL of 100 mM ATP (pH 7.5). The suspension was centrifuged at 4°C for 10 min at 9,000 RCF, and supernatant was removed. The resin was washed twice with 30 mL of Buffer KC (50 mM HEPES-KOH, pH 7.5, 750 mM KCl, 5 mM MgCl_2_, 30 mM imidazole) plus fresh PI and PMSF. Washes consisted of resuspension and hand nutation for 30 sec, 3 min on ice, a 2 min spin at 5,500 RCF in chilled rotors, 3 additional min on ice, followed by supernatant removal. Final supernatant was removed, and the resin was incubated three times with 500 uL of Buffer KE (50 mM HEPES-KOH, pH 7.5, 750 mM KCl, 5 mM MgCl_2_, 300 mM imidazole), centrifuged for 30 sec at 6,000 RCF and supernatant was passed through a filter column to elute proteins. All elutions were then concentrated by Vivaspin column (Cytiva) and placed into dialysis tubing, then incubated overnight with stirring in 1 L of chilled Buffer KL (50 mM HEPES-KOH, pH 7.5, 30 mM KCl, 5 mM MgCl_2_) to remove imidazole. The SUMO protease Ulp1 (lab isolated) was added to the dialyzed protein sample and incubated at room temperature for 1 hr. 500 uL of Buffer KL-equilibrated resin was added, and the mixture was nutated at 4°C for 1 hour. A filter column was used to separate beads from cleaved protein isolate. 10% glycerol was added and proteins were either frozen at −80°C or immediately further purified using an AKTA pure ion exchange chromatography system (Cytiva) and HiTrap Q HP columns (Cytiva), testing fraction activity by quantifying ATP hydrolysis. Proteins were again concentrated with 10% glycerol, separated into aliquots and snap-frozen at −80°C. Isolated Ydj1 was a generous gift from Elizabeth Craig and Sse1 was from a previously isolated laboratory stock ^61^. Purified Hsp104 protein was a kind gift of the Tsai laboratory (Baylor College of Medicine).

### ATP binding assay

For each respective protein sample, 25 uL of ATP-agarose (MilliporeSigma) bead volume was washed three times with 1 mL chaperone buffer (50 mM HEPES-KOH, pH 7.5, 50 mM KCl, 5 mM MgCl_2_, 5 mM DTT) in a siliconized tube. Beads were resuspended in 500 uL chaperone buffer with 1 uM final concentration of protein isolate. Suspensions were nutated at 4°C for 30 minutes and washed five times with 1 mL chaperone buffer + 1.5% Triton X-100 (30 sec spin at 6,000 RCF, on ice in between). After removing supernatant, beads were transferred to a new siliconized tube to negate tube-bound protein and 50 uL of chaperone buffer plus 50 uL of 2X SDS-PAGE sample buffer were added, followed by incubating at 65°C for 20 min prior to gel loading to elute proteins.

### ATPase assay

ATP hydrolysis was determined using a malachite green-based assay (MilliporeSigma) to measure phosphate release. Purified Ssa1 was diluted to 0.1 uM, with or without 0.2 uM Ydj1, in 20 uL total volume of reaction buffer in a 96-well plate. 10 uL of 4mM ATP was added to each well, and the plate was incubated with a cover for 90 min at 30°C, followed by the addition of 150 uL of the malachite green reagent. The colorimetric reaction proceeded for 30 min at room temperature before measuring absorbance. Absorbance values were read and converted to picomoles of phosphate using a standardized phosphate curve. Values are reported as specific activity (pmol ATP/µg Hsp70/min).

### in vitro firefly luciferase recovery assay

Refolding of denatured FFL was assessed as reported previously, with slight alteration ^33^. In short, 200 uM FFL protein was incubated 1:1 with denaturing buffer (50 mM HEPES-KOH, pH 7.5, 50 mM KCl, 5 mM MgCl_2_, 5 mM DTT, 3 M guanidinium HCl) at room temperature for 30 min to denature. Denatured FFL (1 uM) was added to a chaperone mixture containing the respective Ssa1 (0.1 uM), Ydj1 (0.2 uM), Sse1 (0.05 uM), 5 mM ATP, and Hsp104 when applicable (0.1 uM), then brought to a final reaction volume of 100 uL with chaperone buffer. 5 uL of reaction mixture was diluted into a well containing 200 uL of chaperone buffer prior to measurement. To measure activity of properly folded luciferase, 20 uL of luciferin (222 uM) was added to 10 mL of chaperone buffer, and 10 uL was auto-injected into each well and chemiluminescence signal measured. Activity was determined at indicated time points, and raw numbers were converted to a percentage through comparison to activity of a non-denatured control in the same volume.

### Preparation of cell extracts and immunoblotting

Cells were grown overnight, sub-cultured and grown to mid-log phase (OD_600_=0.6-0.8). Proteins were extracted by glass bead lysis as previously described ^62^. Proteins were analyzed by separation on SDS-PAGE gels (8-12%) and transferred to polyvinylidene difluoride (PVDF) membrane (EMD Millipore). Immunoblots were imaged using an anti-Ssa1/2 polyclonal antibody at a 1:10,000 dilution, anti-FLAG mAb at a 1:4,000 dilution (MilliporeSigma), or anti-GFP at a 1:5,000 dilution (Roche) using a previously described procedure _62_. Blots were sprayed with WesternBright ECL-spray (Advansta) and imaged using the ChemiDoc MP Imaging System (Bio-Rad). Bands were quantified using Image Studio Lite (LI-COR Biosciences). To monitor chaperone stability, 100 ug/mL cycloheximide was added to log phase cultures.

### Hsf1 immunoprecipitation

Immunoprecipitation was performed as previously described ^28^. In short, 30 mL of cells were lysed by glass beads, and total lysate was co-incubated with anti-FLAG M2 Affinity gel (MilliporeSigma) in a total volume of 700 uL of TEGN buffer (20 mM Tris-HCl, pH 7.9, 0.5 mM EDTA, 10% glycerol, 50 mM NaCl), plus protease inhibitors (PI), nutating for 2 hr at 4°C. Beads were washed eight times using 750uL of TEGN + PI, followed by elution of proteins using 40 uL of FLAG peptide (200 ug/mL) at room temperature for 25 min. 6X SDS-PAGE sample buffer (350 mM Tris-HCL, pH 6.8, 36% glycerol, 10% SDS, 5% beta-mercaptoethanol, and 0.012% bromophenol blue) was added to samples and incubated at 65°C for 20 min to elute.

### Fluorescence microscopy

For all experiments, yeast live cells were imaged as described previously ^63^. In short, cells were wet-mounted on slides and imaged using the 100x objective of an Olympus IX81 microscope, using a FITC filter to visualize GFP. For each experiment, identical exposure times were used. For foci counts, strains were grown overnight in SC-HIS-URA and late growth stage cells were imaged.

### In vivo FFL assays

FFL-GFP *de novo* assays were carried out as previously specified ^45^. Briefly, indicated strains containing the 413TEF *SSA1-*expressing plasmid and p425MET25-FFL-GFP-leu2::URA3 were grown overnight in SC-URA-HIS at 30°C. 100 uL additional methionine was added to repress plasmid expression. Cells were sub-cultured in fresh media to log phase OD_600_=0.8, 5 mL of cells were harvested by centrifugation, and washed to remove all methionine. Cells were resuspended in 5 mL of SC-URA-MET to induce FFL-GFP expression and activity was measured at the indicated time points by adding 10 uL of 222 nM luciferin in a microplate reader.

Refolding assays were performed similarly to *de novo* FFL activity assays, but after 1 hr of induction, 100 mg/mL of cycloheximide was added to stop protein synthesis. Cells were then subjected to heat denaturing at 42°C for 15 min, then incubated at 30°C for recovery. FFL-GFP activity was measured by adding 10 uL of 222 nM luciferin in a microplate reader or imaged for foci.

### tGND-GFP protein turnover

Steady state levels of the terminally misfolded protein tGND were performed essentially as described ^49^. In brief, BY4741 *ssa1Δssa2Δ* cells containing the pTHD3HA-tGND-GFP plasmid and relevant p413TEF *SSA1* allele plasmid were grown to early log phase in SC-HIS-URA (OD_600_=0.5). 1 mL of cells were collected, washed, resuspended in 177 uL of 1.85 M NaOH and left on ice for 10 min. 177 uL of cold 55% trichloroacetic acid was added, and the sample incubated on ice for an additional 10 min. Cells were centrifuged in a cold room at 7,200 RCF for 1 min and supernatant removed. The pellet was resuspended in 500 uL of ice-cold acetone and centrifuged. The supernatant was removed and 100 uL per OD of 2X urea buffer (1 % SDS, 8 M Urea, 10 mM MOPS, 10 mM EDTA, pH 6.8, 0.01 % bromophenol blue, 1 mM PMSF) was added. Suspensions were sonicated for 5 min and samples placed at 65°C for 20 min, followed by an additional 5 min of sonication. Cells were again centrifuged and supernatant was moved to a new tube and frozen prior to SDS-PAGE separation and immunoblot. For microscopy, cells were grown to early log phase (OD_600_=0.5) in SC-HIS-URA and 100 ug/mL cycloheximide was added to cultures. Cells were imaged for foci at relevant time points.

### Statistical analysis

Student’s *t*-test was used to analyze mean differences between conditions. Prism 9 (GraphPad Software) was used to analyze averages of end point measurements of each time point and calculate standard error of the mean. For all significant tests, \**p*=0.05; \*\**p*=0.005; \*\*\**p*=0.0005; \*\*\*\**p*=0.00005.

## Abbreviations

SBD: substrate-binding domain
NBD: nucleotide-binding domain
FFL: firefly luciferase
HSR: heat shock response
ROS: reactive oxygen species
NEM: *N*-ethylmaleimide

## Acknowledgments

We thank Drs. Qinglian Liu (Virginia Commonwealth University), Randy Hampton (UC San Diego), Nadinath Nillegoda (Monash University), Elizabeth Craig (University of Wisconsin), Andrew Truman (University of North Carolina, Charlotte) and Francis Tsai (Baylor College of Medicine) for advice and reagents. We additionally thank Santosh Kumar and Anna Konavalova (McGovern Medical School) for protein purification assistance. This study was supported by the National Institutes of Health grant R01GM127287.

## Author Contributions

AS and KAM planned and outlined the scope of the work and manuscript. AS performed all the experiments, wrote the initial drafts of the manuscript and was responsible for all revisions. KAM edited the manuscript. AS and KAM designed and created the figures.

