## Supplemental Information for "Redox regulation of yeast Hsp70 modulates protein quality control while directly triggering an Hsf1-dependent cytoprotective response"

### Santiago and Morano
### Supplementary Information

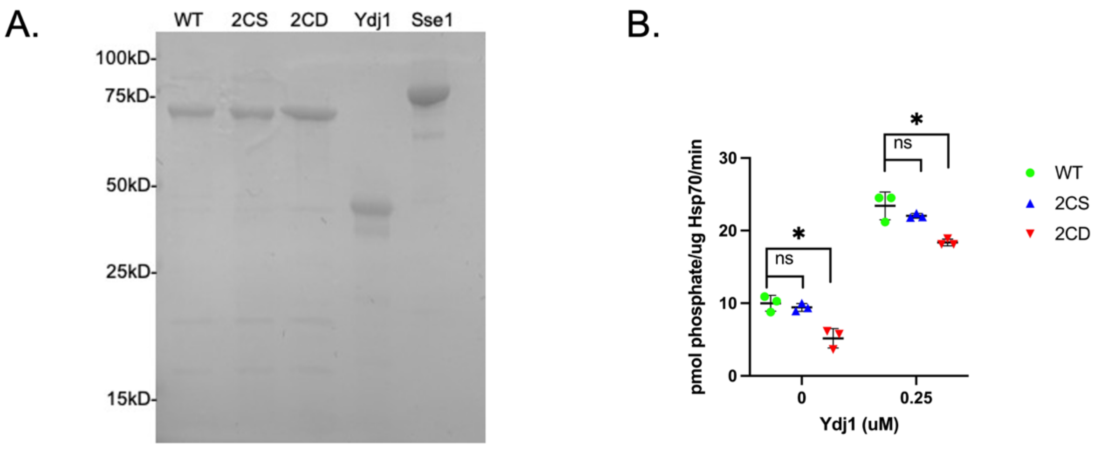


**Fig. S1. The oxidomimetic Ssa1-2CD mutant exhibits reduced basal and Ydj1-stimulated ATPase activity.** (A) Coomassie staining of approximately equivalent amounts of the indicated proteins used in the *in vitro* assays. (B) ATP hydrolysis by wild type and mutant Ssa1 proteins, basal and stimulated by 0.2 µM Ydj1. Bolded horizontal bars indicate mean, and error bars indicate SEM. *, $p<0.05$; ns, not significant.

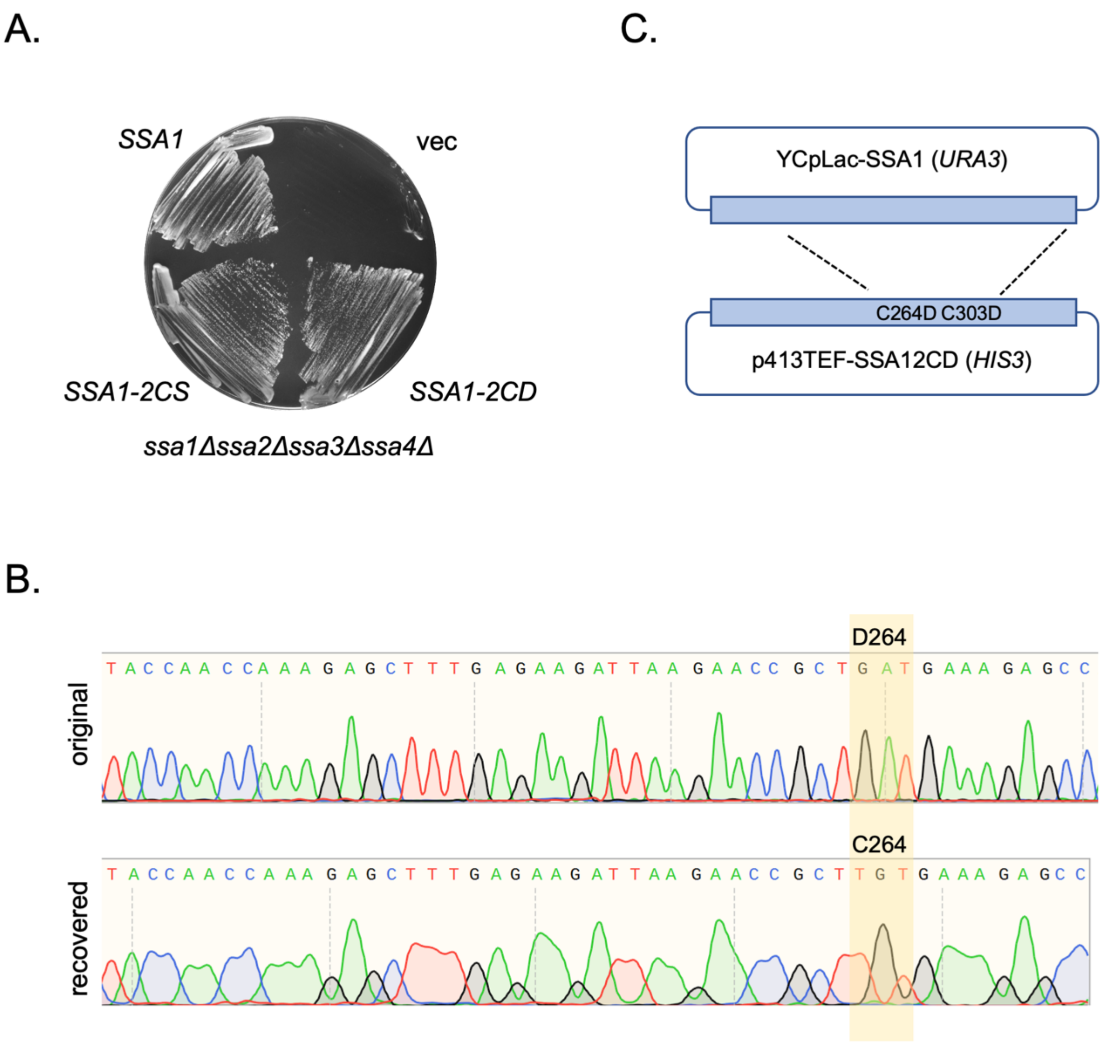


**Fig. S2. The oxidomimetic *ssa1-2CD* mutant is incapable of supporting viability as the sole cytosolic *SSA* gene.** (A) 48-hour 5-fluoroorotic acid (5-FOA) plate growth of the indicated *HIS3*-selective plasmid, in an *ssa1Δ ssa2Δ ssa3Δ ssa4Δ* deletion background demonstrating unexpected wild type growth for the *ssa1-2CD* mutant. (B) Sequencing analysis of the introduced *HIS3*-selective plasmid (original) and the recovered plasmid extracted from the colonies shown in (A), demonstrating reversion of the aspartic acid-encoding codon GAT to the original cysteine-encoding codon TGT. Only the region surrounding C264 is shown for simplicity. (C) Proposed recombination mechanism to explain the gene reversion event during the plasmid shuffle process.

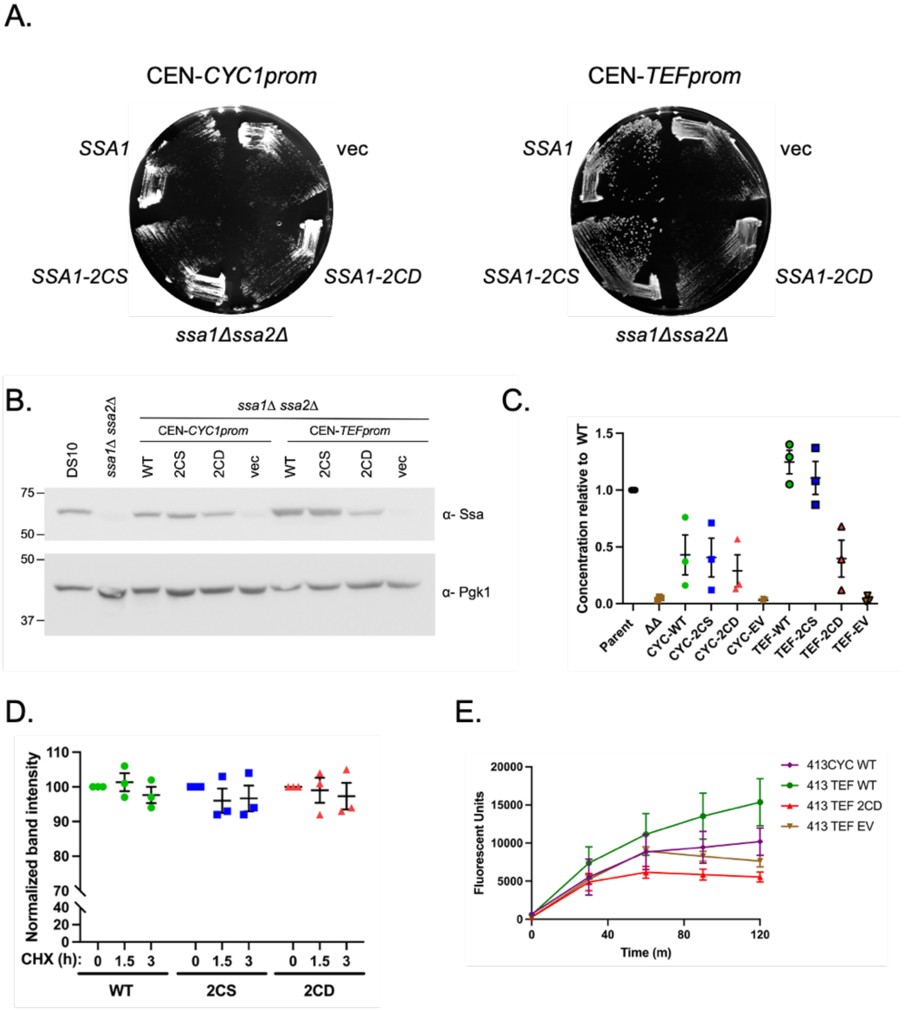


**Fig. S3. Expression of the Ssa1-2CD protein is restricted relative to wild type Ssa1 and Ssa1-2CS**. (A) 48-hour plate growth of each indicated *SSA* gene driven from the weak *CYC1* promoter and the stronger *TEF* promoter in the *ssa1Δ ssa2Δ* background. (B) Relative protein expression of each indicated Ssa1 protein from the wild type strain DS10, or the respective *CYC1* or *TEF* plasmid in the *ssa1Δ ssa2Δ* background. (C) Quantification of the relative levels of each indicated protein compared to expression in the DS10 background. (D) Immunoblot cycloheximide chase analysis to monitor protein stability over time of each indicated protein driven from the *TEF* promoter, in the *ssa1Δ ssa2Δ* background. (E) *de novo* expression of FFL-GFP. Bolded horizontal bars indicate mean, and error bars indicate SEM.
